# *Pseudomonas aeruginosa* increases the susceptibility of *Candida albicans* to amphotericin B in dual species biofilms

**DOI:** 10.1101/2022.10.19.512978

**Authors:** Farhana Alam, Sarah Blackburn, Joao Correia, Jessica Blair, Rebecca A Hall

## Abstract

Biofilms are the leading cause of nosocomial infections, and are hard to eradicate due to their inherent antimicrobial resistance. *Candida albicans* is the leading cause of nosocomial fungal infections, and is frequently co-isolated with the bacterium *Pseudomonas aeruginosa* from biofilms in the Cystic Fibrosis lung and severe burn wounds. The presence of *C. albicans* in multi-species biofilms is associated with enhanced antibacterial resistance, which is largely mediated through fungal extracellular carbohydrates sequestering the antibiotics. However, significantly less is known regarding the impact of polymicrobial biofilms on antifungal resistance. Here we show that, in dual species biofilms, *P. aeruginosa* enhances the sensitivity of *C. albicans* to amphotericin B, an effect that was biofilm specific. Transcriptional analysis combined with gene ontology enrichment analysis identified several *C. albicans* processes associated with oxidative stress to be differentially regulated in dual species biofilms, suggesting that *P. aeruginosa* exerts oxidative stress on *C. albicans*, likely through the actions of secreted phenazines. However, the mitochondrial superoxide dismutase *SOD2* was significantly downregulated in the presence of *P. aeruginosa.* Monospecies biofilms of the *sod2*Δ mutant were more susceptible to amphotericin B, and the susceptibility of these biofilms was further enhanced by the addition of exogenous phenazines. Therefore, we propose that in dual species biofilms, *P. aeruginosa* simultaneously induces mitochondrial oxidative stress, whilst downregulating key detoxification enzymes, which prevent *C. albicans* mounting an appropriate oxidative stress response to amphotericin B, leading to fungal cell death. This work highlights the importance of understanding the impact of polymicrobial interactions on antimicrobial susceptibility.

**Importance:** Biofilms are aggregates of cells encased in an extracellular matrix, and are responsible for the majority of infections in hospitals. The Gram-negative bacterium *Pseudomonas aeruginosa*, and the fungal pathogen *Candida albicans* are frequently co-isolated from biofilms in the Cystic Fibrosis lung, and in burn wounds. Here we show that in these biofilms, *P. aeruginosa* secreted phenazines induce mitochondrial reactive oxygen species stress, which enhances the sensitivity of *C. albicans* to the antifungal amphotericin B. This work highlights the importance of understanding the impact of polymicrobial interactions on antimicrobial susceptibility.

## Introduction

*Candida albicans* is a commensal and opportunistic fungal pathogen of humans that is frequently co-isolated from infection sites with the Gram-negative bacterium *Pseudomonas aeruginosa* (Dhamgaye *et al.* 2016). Extensive research has shown that these two microbes undergo a complex range of interactions, the outcome of which appears to be dependent on the surrounding environment. For example, *P. aeruginosa* has been shown *in vitro* to bind and kill *C. albicans* hyphae (Hogan and Kolter 2002), while *in vivo, P. aeruginosa* and *C. albicans* have been shown to have an agonistic relationship, resulting in enhanced pathogenesis (Bergeron *et al.* 2017). *P. aeruginosa* secretes a broad range of virulence factors including phenazines, siderophores, haemolysins, and phosphatases, which are regulated by one or more quorum sensing systems. Several of these secreted products have been shown to directly affect *C. albicans*, with the phenazines resulting in the induction of reactive oxygen species (ROS) induced stress, and inhibition of fungal growth and morphogenesis (Morales *et al.* 2013, Tupe *et al.* 2015), while siderophores impose nutrient restriction. On the other hand, *C. albicans* secretes the quorum sensing molecule farnesol, which inhibits the *Pseudomonas* Quinolone Signal (PQS) pathway (Cugini *et al.* 2007), and imposes ROS induced stress on other microbes (Machida et al. 1998, Liu et al. 2010).

*C. albicans* and *P. aeruginosa* are most commonly co-isolated from the lungs of Cystic Fibrosis patients (Doern and Brogden-Torres 1992), and from wounds of burn victims (Weaver *et al.* 2019), where the two microbes tend to grow together in biofilms. Biofilms are communities of cells encased in an extracellular matrix (ECM), which provides protection from the host’s immune responses, and increases the resistance of microbes to antimicrobial agents. Cells in a mono-species biofilm can exhibit MIC values 100–1000-fold greater than their planktonic counterparts. However, in mixed-species biofilms, the microbes interact through secreted signalling molecules, direct cell-cell interactions and through nutrient competition (Fourie and Pohl 2019), all of which have the potential to affect antimicrobial resistance. Furthermore, each species contributes specific extracellular polysaccharides to the ECM, and fungi produce hyphae which act as molecular scaffolds that together alter the structure and composition of the biofilm. These interactions affect the sensitivity of the microbes to antimicrobial therapy. For example, the presence of *C. albicans* in bacterial biofilms promotes the resistance of *S. aureus, P. aeruginosa* and *E. coli* to a variety of antibiotics (Harriott and Noverr 2009, De Brucker *et al.* 2015, Kong *et al.* 2016, Alam *et al.* 2020). Although the precise molecular mechanisms behind this enhanced antibacterial resistance are not fully understood, it is thought that the contribution of fungal extracellular polysaccharides to the ECM sequesters the antibiotic molecules and limits their diffusion through the biofilm. Furthermore, microbe-microbe and nutrient competition may alter the transcriptional profile of bacterial cells, priming the cells to be more tolerant to the antibiotic. However, how these microbe-microbe interactions affect the response to antifungal treatment is unknown.

Currently there are only a handful of classes of antifungal drugs on the market, the most widely used of which are the azoles (fluconazole, miconazole, etc.), which target ergosterol biosynthesis, the echinocandins (caspofungin and anidulafungin) that target the synthesis of beta-glucan, and the polyenes (amphotericin B and nystatin) that target ergosterol. The widespread use of antifungals in both hospital and agricultural settings has resulted in the rapid rise of antifungal resistance, with the WHO identifying multidrug-resistant *Candida* infections as a major cause for concern. With limited new antifungal drugs in the pipeline, there is an urgent need to preserve and improve the efficacy of the available antifungals. Here, we demonstrate that in a dual-species biofilm *P. aeruginosa* enhances the sensitivity of *C. albicans* to the antifungal amphotericin B, but has minimal effect on the sensitivity of the fungus to fluconazole or caspofungin. The enhanced sensitivity of *C. albicans* to amphotericin B was mediated by the *P. aeruginosa*-dependent repression of the fungal superoxide dismutase, *SOD2,* resulting in *C. albicans* being exposed to enhanced reactive oxygen stress.

## Methods

### Strains and Media

*C. albicans* strains were maintained on YPD agar and grown in YPD broth, while *P. aeruginosa* strains were grown and maintained on LB medium. All biofilm assays were performed in Mueller-Hinton broth (MHB). The strains used in this study are listed in Table S1.

### Biofilm assay

Biofilm assays were performed as described previously (Alam *et al.* 2020). In brief, overnight cultures of *C. albicans* and *P. aeruginosa* were washed in PBS, and *C. albicans* resuspended at 1×10^6^ cells/ml and *P. aeruginosa* to OD600 of 0.2 in MHB. Each well of 96-well plates contained 100 μl *C. albicans* and 10 μl of *P. aeruginosa.* Plates were incubated at 37°C for 2 hrs to allow cells to adhere, at which point the media was replaced with fresh sterile media, and plates incubated statically at 37°C for 24 hrs. Cells not part of the biofilm were removed and media replaced with fresh MHB containing the appropriate concentration of antifungal agent, and plates incubated for 2 hrs. Media was replaced with 100 μl PBS containing 50 μg/ml DNase I and plates incubated at 37°C for 1 hr to degrade the extracellular matrix. Biofilms were detached from the plates by scraping, serially diluted, and plated onto YPD agar supplemented with 100 μg/ml tetracycline to determine viable *C. albicans* CFUs.

To assess the effects of phenazines or H_2_O_2_ on antifungal resistance, compounds were added to single species biofilms at the indicated concentrations, after the adherence step, and biofilm formed as outlined above.

### RNA sequencing

Biofilms were formed as described previously (Alam *et al.* submitted). Triplicate biofilms were pooled, and 50 μl serially diluted and plated on cetrimide agar or YPD supplemented with 100 μg/ml tetracycline to check for contamination. Remaining biofilm cells were centrifuged at 3500 rpm at 4°C for 5 min, and pellets snap frozen in liquid nitrogen. Four biological replicates were shipped to GeneWiz®, UK, for RNA extraction, sequencing and basic bioinformatic analysis.

### RNA extraction and sequencing

RNA was extracted from the biofilms by GeneWiz®using the Qiagen RNeasy Plus mini kit. Library preparation was done in the following stages: A) ribosomal RNA depletion; B) RNA fragmentation and random priming; C) first and second strand cDNA synthesis; D) end repair, 5’ phosphorylation and dA-tailing; E) adapter ligation, PCR enrichment and sequencing. Paired-end sequencing was performed using Illumina HiSeq 4000 (2×150bp configuration, single index, per lane).

### Bioinformatic analysis

Sequence quality of each sample was evaluated by determination of the number of reads, the yield (Mbases), the mean quality score, and the percentage of reads over 30 bases in length. FastQC software was used to determine per base sequence quality and per sequence GC content. Sequence reads were trimmed to remove adapter sequences and nucleotides with poor quality, using Trimmomatic v.0.36. The trimmed reads were mapped to the *C. albicans* reference genome, available on ENSEMBL, using the STAR aligner v.2.5.2b. For dual species biofilms, reads were first mapped to the *P. aeruginosa* PAO1 genome available on ENSEMBL, and then unmapped reads were aligned to the *C. albicans* genome. Unique gene hit counts were calculated using featureCounts from the Subread package v.1.5.2. Only unique reads that fell within exonic regions were counted (the maximum hits per read was set to 10 by default). One biological replicate of the *C. albicans* mono-species biofilms (CA 0M-6) had a significant number of reads that did not align to the *C. albicans* genome or the *P. aeruginosa* genome and was therefore excluded from downstream analysis. Differential gene expression analysis was performed using DESeq2. Principal component analysis (PCA) was performed to reveal the similarities within and between groups, with PCA plots included in the output (Fig S1). The DESeq2 output file (*C. albicans*_expression_CA_vs_PACA.xlsx) and the raw and normalise reads and TPM values for all *C. albicans* (*C_albicans*_counts.xlsx) and *P. aeruginosa* (*P_aeruginosa*_count_data.xlsx) genes are available at the Gene Expression Omnibus (GEO) database (https://www.ncbi.nlm.nih.gov/geo/), under accession number GSE167137.

### Enrichment analysis

For *C. albicans* transcriptomic analysis, differential expression of genes between conditions was considered significant if the adjusted P-value (Padj) was ≤0.05 and the log_2_-fold change was ≥1 (Bruno *et al.* 2015, Dutton *et al.* 2016, Fourie *et al.* 2021). Gene ontology (GO) analysis was performed using the *Candida* Genome Database (CGD) Gene Ontology Slim Mapper (Arnaud *et al.* 2009, Skrzypek *et al.* 2017). This tool maps the annotations from each gene list to GO Slim terms, which are broad, high level GO terms that are specific to the selected *Candida* species; in this case, *C. albicans* (Skrzypek *et al.* 2017). KEGG pathway enrichment analysis was done using KOBAS 3.0 software (Xie *et al.* 2011), as described previously (Alam *et al.* submitted).

### Quantification of ergosterol

Biofilms were formed as described above. After the 24-hr incubation, biofilms were washed with PBS and fixed with 4% PFA for 1 hr at room temperature. Biofilms were disrupted and then stained with filipin (0.05 mg/ml) for 2 hrs at room temperature. Stained biofilm cells were washed with PBS, mounted onto glass microscope slides, and imaged on a Zeiss AxioObserver at 63x magnification, acquiring 30-33 images per sample. To quantify the amount of ergosterol, the median fluorescent intensity of the filipin staining was quantified using FIJI software. Up to five cells per image were outlined, as well as five spherical areas containing no cells (background fluorescence readings). The mean corrected total cell fluorescence (CTCF) was calculated for each image, in which CTCF = integrated density - (area of selected cell x mean fluorescence of background readings). Data were analysed using one-way ANOVA and Holm-Sidak’s multiple comparisons test.

## Results

### *P. aeruginosa* increases the susceptibility of *C. albicans* to amphotericin B

Previously we have shown that dual-species biofilms of *C. albicans* and *P. aeruginosa* have increased tolerance to antibiotics, as a result of the fungal extracellular matrix sequestering the drugs (Alam *et al.* 2020). Therefore, we hypothesised that *P. aeruginosa* may also impact the antifungal susceptibility of *C. albicans*. To test this hypothesis, 24-hr mono- or dual-species biofilms were treated with antifungal drugs for either 18 (fluconazole and caspofungin) or 2 (amphotericin B) hours and then *C. albicans* viability quantified by plating. The susceptibility of *C. albicans* in dual-species biofilms to fluconazole was not affected compared with the mono-species control, with significant fungal survival observed when biofilms were treated with high concentrations of the azole (1000 μg/ml) (Fig 1A). *P. aeruginosa* had a slight impact on the sensitivity of *C. albicans* to caspofungin (Fig 1B), with *C. albicans* survival being reduced at high concentrations (0.5–1 μg/ml). However, treatment of the dual-species biofilms with the polyene amphotericin B, resulted in a significant reduction in *C. albicans* viability, with negligible fungal viability detected when the dual-species biofilms were treated with 2 μg/ml amphotericin B (Fig 1C). On the other hand, treatment of mono-species biofilms with amphotericin B still resulted in significant fungal survival, even when treated with 8 μg/ml amphotericin B. Therefore, these data suggest that the presence of *P. aeruginosa* in the biofilm results in a significant increase in the susceptibility of *C. albicans* to the polyene class of antifungals.

**Figure 1.**
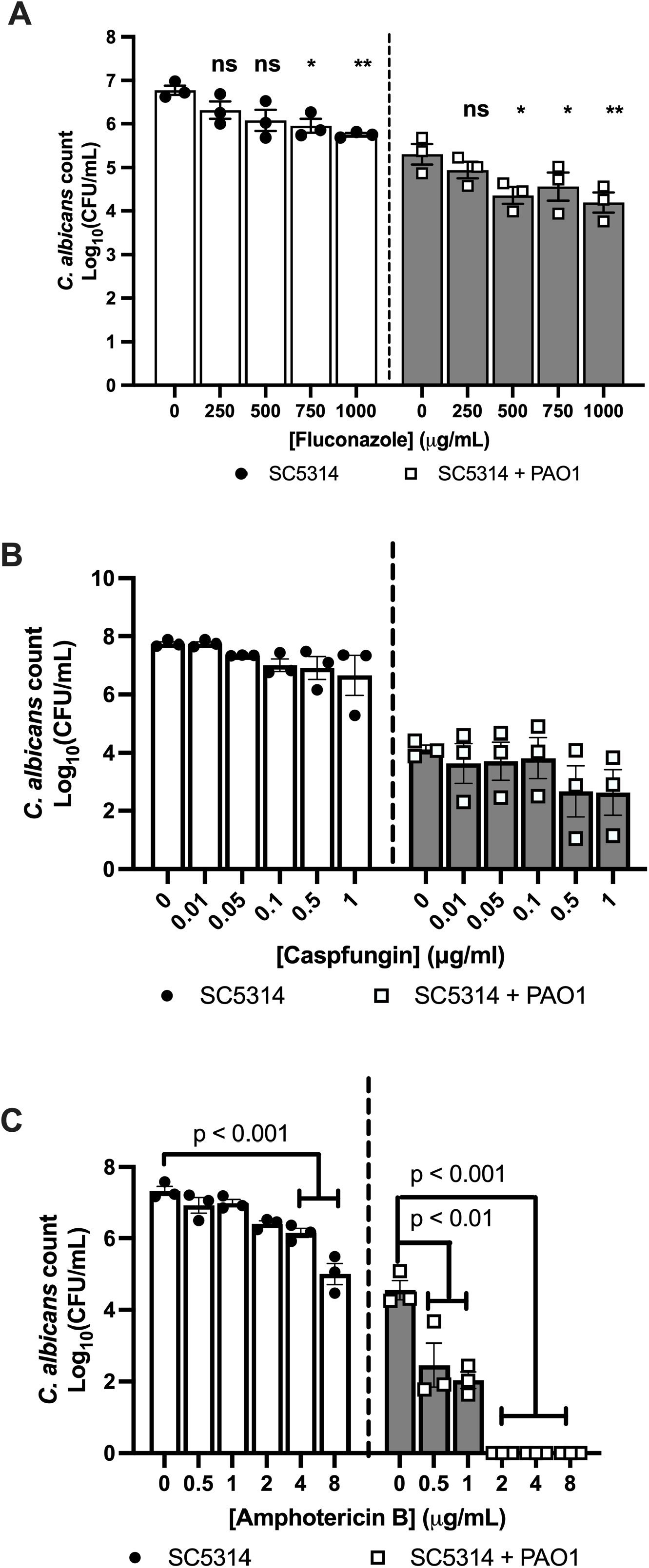
*P. aeruginosa* enhances the sensitivity of *C. albicans* to amphotericin B in dual-species biofilms. 24-hr preformed single- or dual-species biofilms were treated with increasing concentrations of **A)** fluconazole, **B)** caspofungin or **C)** amphotericin B for 18 (A and B) or 2 hrs (C). Data are the log_10_(mean) +/− the SEM from 3 biological replicates. Data were analysed using 2-way ANOVA and Holm-Sidak’s multiple comparisons test.

To determine the role of *P. aeruginosa* in this interaction, we grew *C. albicans* biofilms in the presence of heat-killed or fixed *P. aeruginosa* cells, and added live *P. aeruginosa* to mature *C. albicans* mono-species biofilms just prior to the addition of amphotericin B. *C. albicans* susceptibility to the polyene was only increased in the presence of live bacteria, which had been directly incorporated into the *C. albicans* biofilm (Fig 2A, B), suggesting that bacterial viability and the architecture of the biofilm are important for the observed phenotype. In agreement with this, the susceptibility of *C. albicans* to amphotericin B was not altered when *C. albicans* was co-cultured with *P. aeruginosa* under planktonic conditions (Fig 2C). To confirm that the impact of *P. aeruginosa* on *C. albicans* amphotericin B susceptibility is a general trait of *P. aeruginosa*, we screened several *P. aeruginosa* clinical isolates for their ability to suppress the resistance of *C. albicans* to the polyene. The viability of *C. albicans* was significantly reduced in all dual-species biofilms treated with amphotericin B, although not as significantly as PAO1 (Fig 2D). However, increasing the incubation time of the biofilms with the antifungal from 2 hrs to 18 hrs killed all fungal cells (Fig 2E). Therefore, *P. aeruginosa* enhances the susceptibility of *C. albicans* to amphotericin B in biofilms.

**Figure 2.**
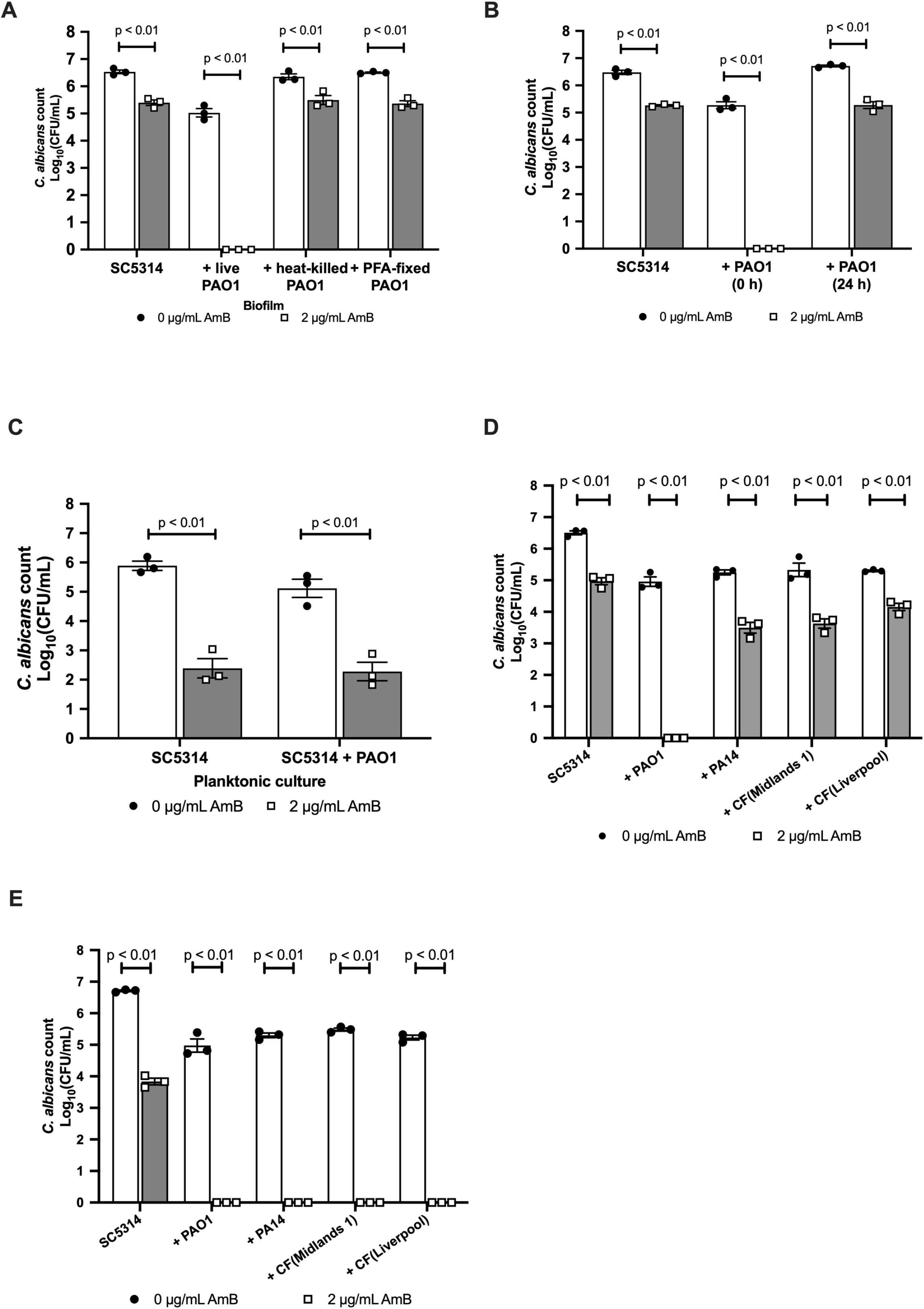
Impact of *P. aeruginosa* on amphotericin B sensitivity is biofilm specific. **A)** SC5314 biofilms were grown in the presence of either heat-killed or PFA-fixed *P. aeruginosa* cells for 24 hrs and then treated with 2 μg/ml amphotericin B for 2 hrs. **B)** SC5314 biofilms were formed with PAO1 cells added at the start (0 hr) of biofilm formation or after *C. albicans* had formed a robust biofilm (24 hrs). **C)** SC5314 was grown planktonically in the presence or absence of PAO1 for 24 hrs and then treated with 2 μg/ml amphotericin B for 2 hrs. **D)** SC5314 biofilms were formed in the presence of different strains of *P. aeruginosa*, including two clinical isolates from Cystic Fibrosis patients, and treated with 2 μg/ml amphotericin B for 2 hrs. **E)** SC5314 biofilms were formed in the presence of different strains of *P. aeruginosa*, including two clinical isolates from Cystic Fibrosis patients, and treated with 2 μg/ml amphotericin B for 18 hrs. Data are the log_10_(mean) +/− the SEM from 3 biological replicates. Data were analysed using 2-way ANOVA and Holm-Sidak’s multiple comparisons test.

### *P. aeruginosa* also increases the susceptibility of *C. dubliniensis* to Amphotericin B

To determine whether *P. aeruginosa* impacts the susceptibility of other *Candida* species to amphotericin B, we grew *P. aeruginosa* in dual-species biofilms with non-*albicans Candida* species for 24 hrs, and then treated the biofilms with 2 μg/ml amphotericin B. *C. dubliniensis* behaved similarly to *C. albicans*, being more susceptible to amphotericin B in the presence of *P. aeruginosa.* However, *C. tropicalis, C. parapsilosis, C. krusei,* and *C. glabrata* still displayed significant viability after treatment with amphotericin B in both mono- and dual-species biofilms (Fig 3). The viability of *C. tropicalis* was significantly reduced in the presence of *P. aeruginosa*, suggesting that the bacterium has some impact on *C. tropicalis*. On the other hand, the viability of the other *Candida* species in dual-species biofilms was comparable to mono-species biofilms, indicating that *P. aeruginosa* has negligible impact on these *Candida* species.

**Figure 3.**
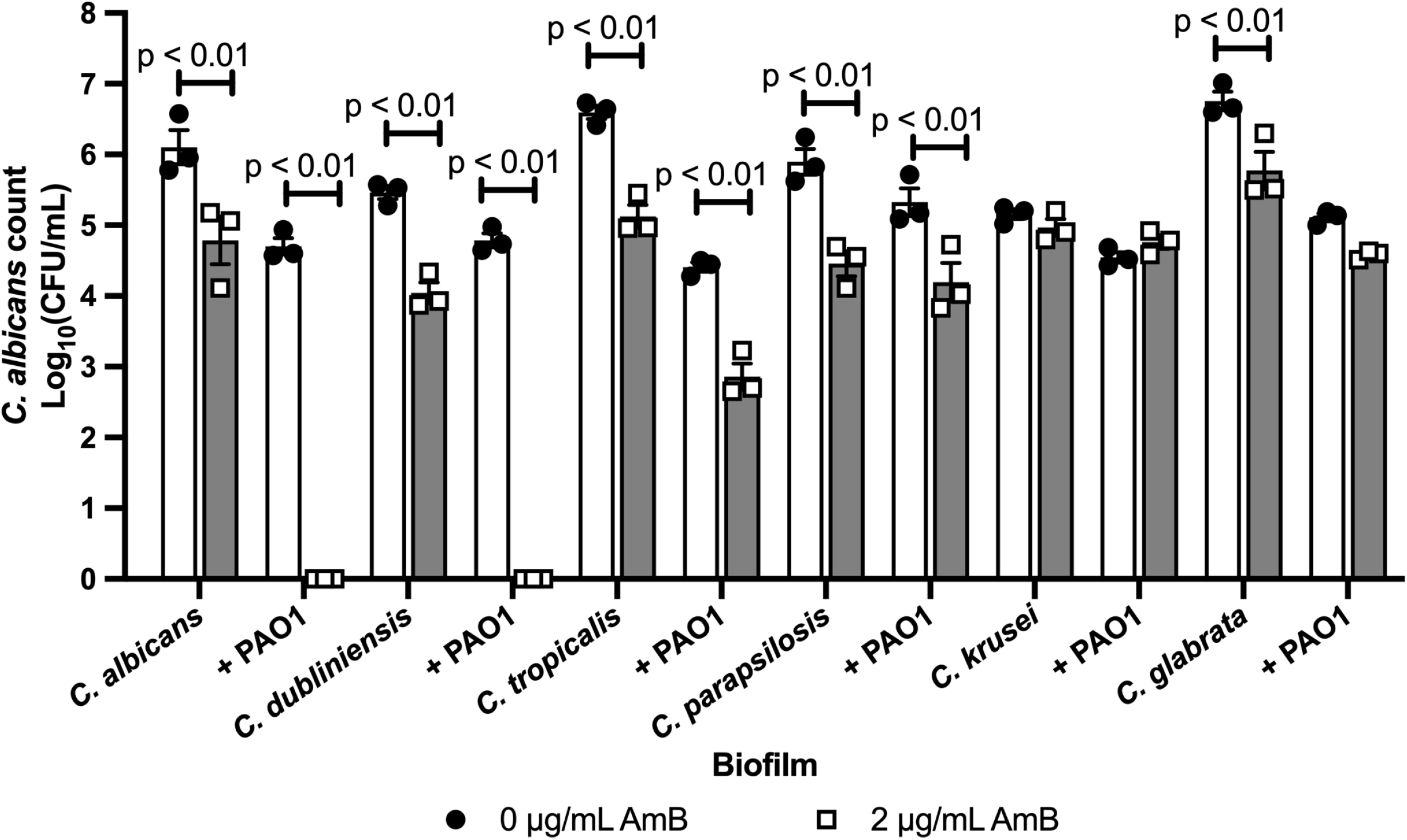
*P. aeruginosa* also increases the sensitivity of *C. dubliniensis* to amphotericin B. Medically important *Candida* species were grown in single- or dual-species biofilms with PAO1 for 24 hrs and then biofilms were treated with 2 μg/ml amphotericin B for 2 hrs. Data are the log_10_(mean) +/− the SEM from 3 biological replicates. Data were analysed using 2-way ANOVA and Holm-Sidak’s multiple comparisons test.

### *C. albicans* mutants with reduced ergosterol content are more resistant to amphotericin B

One of the main modes of action of amphotericin B is to bind ergosterol in the fungal membrane, forming pores resulting in fungal cell lysis. Genes involved in the synthesis of ergosterol are regulated by the Upc2 transcription factor (Silver *et al.* 2004). Therefore, the *upc2Δ* mutant synthesises significantly less ergosterol than wild type cells (Silver *et al.* 2004). To determine whether the increased susceptibility of *C. albicans* to amphotericin in the presence of *P. aeruginosa* is dependent on ergosterol, we quantified the susceptibility of the *upc2Δ* mutant to amphotericin B in both mono- and dual-species biofilms. Deletion of *UPC2* attenuated the ability of *P. aeruginosa* to impact the susceptibility of *C. albicans* to the polyene, with significant fungal viability observed in dual-species biofilms treated with amphotericin B (Fig 4A). Therefore, we hypothesised that *P. aeruginosa* increases the susceptibility of *C. albicans* to the polyene by increasing the amount of ergosterol in the fungal cell membrane. However, the levels of ergosterol were similar between mono- and dual-species biofilms (Fig 4B), suggesting that although ergosterol levels are important for the mode of action of amphotericin B, *P. aeruginosa* is not directly increasing ergosterol incorporation into the fungal cell membrane.

**Figure 4.**
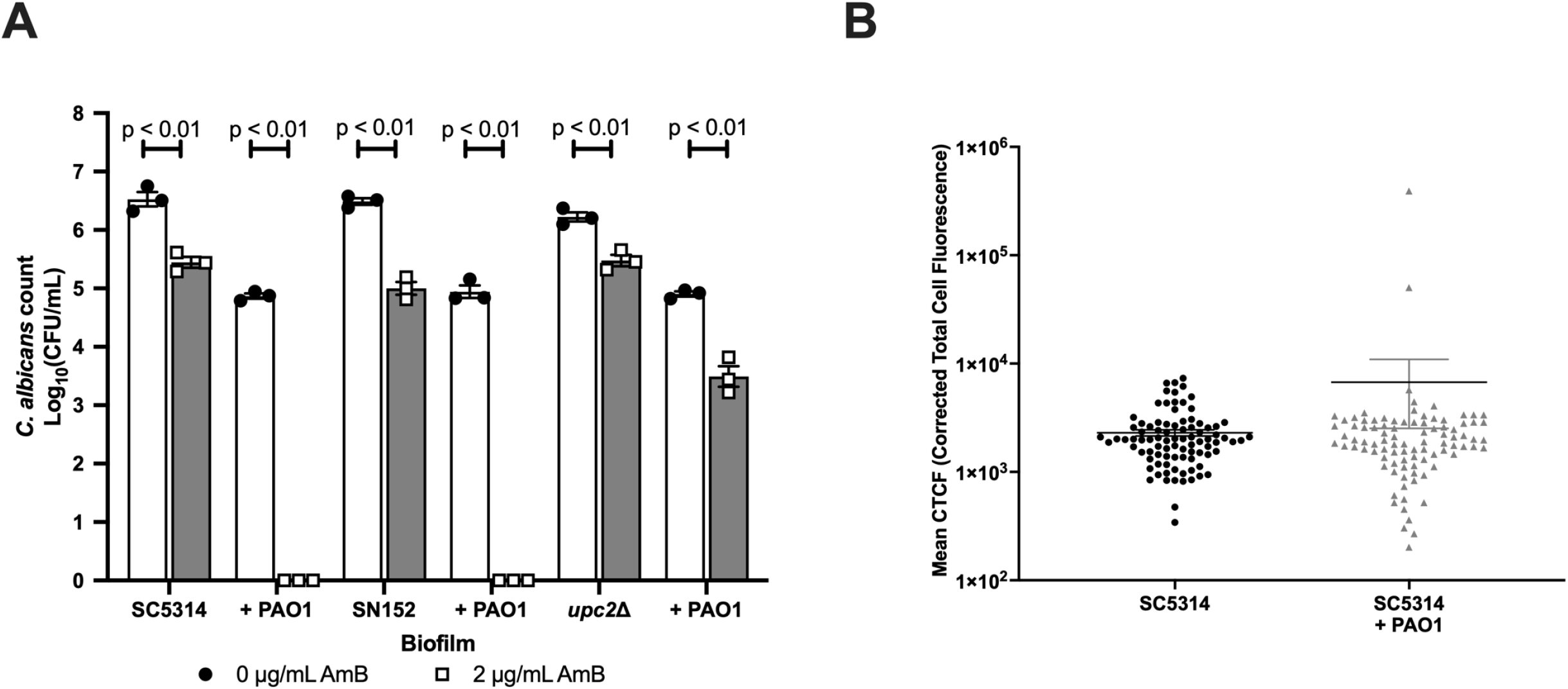
Deletion of *UPC2,* the key transcription factor regulating ergosterol biosynthesis, enhances the resistance of *C. albicans* to amphotericin B in the presence of *P. aeruginosa*. **A)** *C. albicans* strains were grown in single- or dual-species biofilms for 24 hrs and then biofilms were treated with 2 μg/ml amphotericin B for 2 hrs. Data are the log_10_(mean) +/− the SEM from 3 biological replicates. Data were analysed using 2-way ANOVA and Holm-Sidak’s multiple comparisons test. **B)**24-hr preformed biofilms were fixed with PFA, disrupted, and stained with filipin.

### *C. albicans* undergoes a significant transcriptional response to the presence of *P. aeruginosa*

To elucidate how *P. aeruginosa* is increasing the susceptibility of *C. albicans* to amphotericin B, global transcriptional analysis was performed on *C. albicans* in both mono- and dual-species biofilms. The presence of *P. aeruginosa* resulted in a significant transcriptional response in *C. albicans* with 431 genes being upregulated and 739 genes being downregulated (Padj < 0.05, and log_2_-fold cut-off > 1) (Fig 5A). In agreement with our ergosterol staining data, KEGG pathway enrichment analysis identified ergosterol biosynthesis to be among the biological processes that are significantly downregulated in the dual-species biofilms (Fig 5B). Therefore, the increased sensitivity of *C. albicans* to amphotericin B in the presence of *P. aeruginosa* is not due to increased ergosterol levels. Genes associated with filamentous growth and cell wall biosynthesis were also downregulated in dual species biofilms. *P. aeruginosa* binds to *C. albicans* hyphae via O-mannans resulting in fungal cell death (Hogan and Kolter 2002, Brand *et al.* 2008). Therefore, the downregulation of these processes might prevent bacterial attachment and ensure fungal survival. Other processes that were significantly downregulated included genes involved in ribosome biogenesis, translation and carbon metabolism, suggesting that *C. albicans* is under stress in the dual-species biofilms. In agreement with this, genes involved in iron acquisition were significantly upregulated, suggesting that in dual species biofilms there is nutrient competition. Other processes that were upregulated included amino acid metabolism and peroxisome function (Fig 5C). In addition to functioning in fatty acid oxidation, the fungal peroxisome is involved in detoxifying hydrogen peroxide, suggesting that *P. aeruginosa* may be imposing reactive oxygen stress on *C. albicans*. Taken together these results suggest that *C. albicans* is under significant stress, likely reactive oxygen stress, in the dual-species biofilms, resulting in the reduction of protein synthesis.

**Figure 5.**
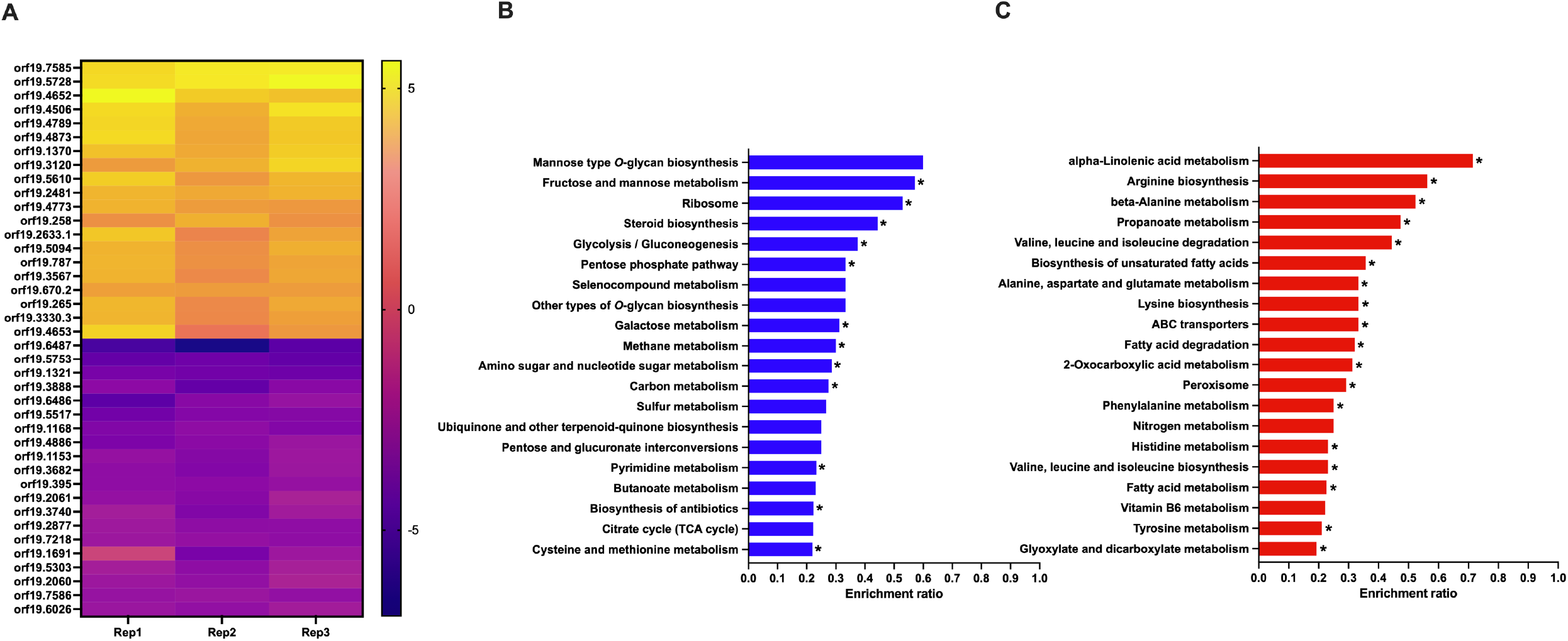
Transcriptional profiling confirms that *C. albicans* is under oxidative stress in dual-species biofilms. **A)** Expression (log_2_) of the 40 most significantly differentially regulated genes. **B)** KEGG enrichment analysis of pathways significantly downregulated in dual-species biofilms. **C)** KEGG enrichment analysis of pathways significantly upregulated in dual-species biofilms.

### *P. aeruginosa* increases the susceptibility of *C. albicans* to amphotericin B through the phenazine-mediated induction of reactive oxygen stress

In addition to binding ergosterol and inducing cell lysis, amphotericin B has been shown to exert antifungal activity through the generation of reactive oxygen species (ROS) (Guirao-Abad *et al.* 2017). At the same time, *P. aeruginosa* is also known to induce ROS stress in *C. albicans* (Tupe et al. 2015). Given that our transcriptomics data suggested that *C. albicans* was under ROS stress in the dual-species biofilms, we hypothesised that ROS induction by both *P. aeruginosa* and amphotericin B at the same time may result in enhanced antifungal activity. In support of this hypothesis, treatment of *C. albicans* mono-species biofilms with hydrogen peroxide resulted in enhanced susceptibility of the fungus to amphotericin B, similar to the effects observed in the presence of *P. aeruginosa* (Fig 6A). Similar effects were observed when *C. albicans* mono-species biofilms were established in the presence of menadione, an inducer of mitochondrial ROS. However, *C. albicans* biofilms grown in the presence of hypoxanthine and xanthine oxidase which induce cellular ROS did not increase the susceptibility of *C. albicans* to amphotericin B (Fig 6A). Therefore, *P. aeruginosa* increases *C. albicans* susceptibility to the polyene through the induction of mitochondrial ROS.

**Figure 6.**
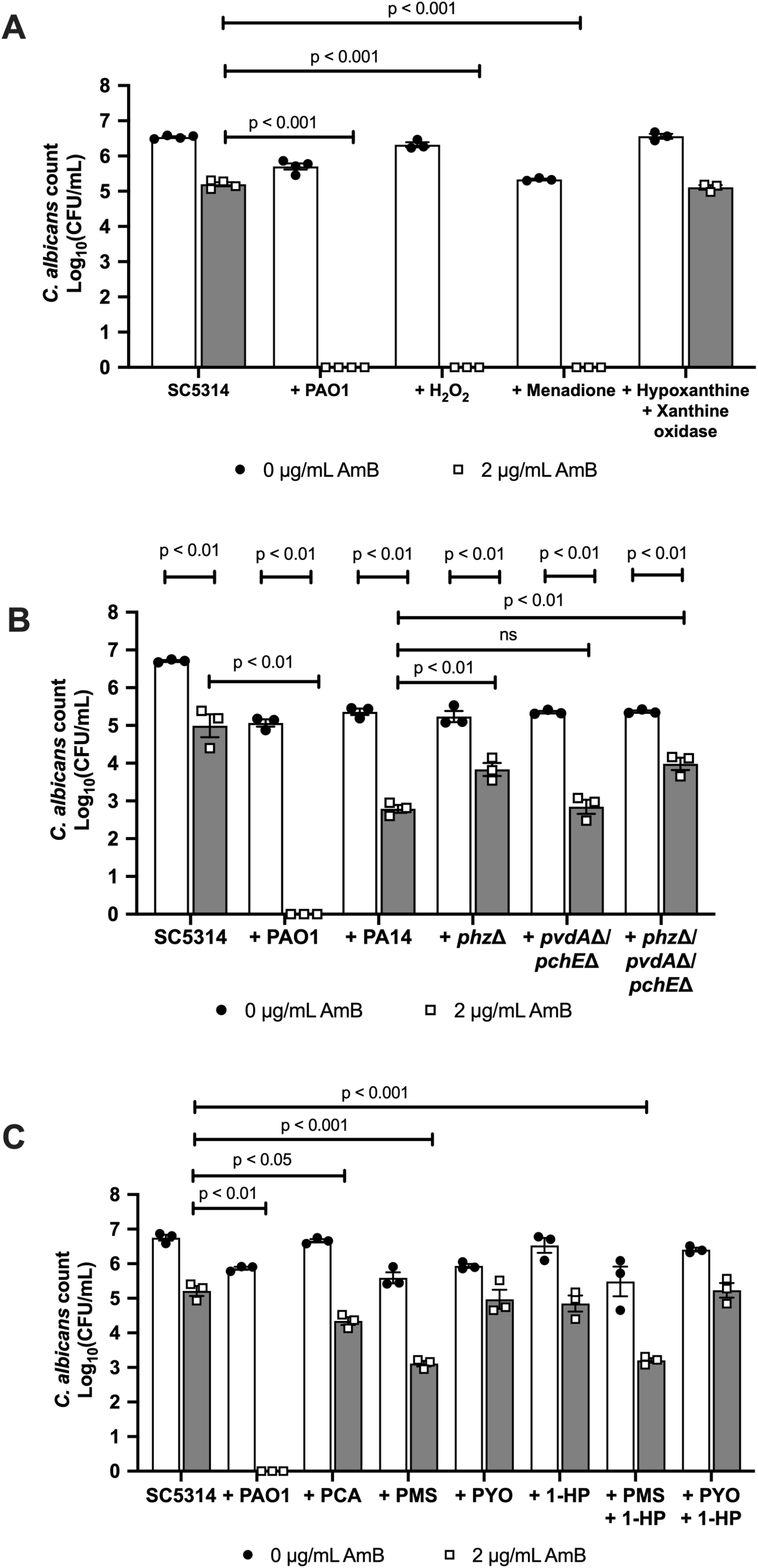
*P. aeruginosa* increased the sensitivity of *C. albicans* to amphotericin B through the induction of mitochondrial ROS. **A)** SC5314 single-species biofilms were grown in the presence of H_2_O_2_, menadione, and hypoxanthine and xanthine oxidase for 24 hrs, and then treated with 2 μg/ml amphotericin B for 2 hrs. **B)** SC5314 was grown in single- or dual-species biofilms with *P. aeruginosa* strains defective in iron and phenazine biosynthesis. **C)** SC5314 single-species biofilms were grown in the presence of purified phenazine compounds for 24 hrs, and then biofilms were treated with 2 μg/ml amphotericin B for 2 hrs. Data are the log_10_(mean) +/− the SEM from 3 biological replicates. Data were analysed using 2-way ANOVA and Holm-Sidak’s multiple comparisons test.

*P. aeruginosa* has been shown to induce ROS stress in *C. albicans* through the secretion of phenazines (Tupe *et al.* 2015). In agreement with phenazine-mediated ROS stress in *C. albicans,* deletion of genes involved in phenazine biosynthesis attenuated the ability of *P. aeruginosa* to enhance the susceptibility of *C. albicans* to amphotericin B (Fig 6B). Addition of exogenous phenazines to *C. albicans* mono-species biofilms confirmed that phenazine methosulfate (PMS), and to a lesser extent phenazine-1-carboxylic acid (PCA), mediated ROS stress and therefore enhanced *C. albicans* sensitivity to amphotericin B (Fig 6C).

### *P. aeruginosa* increases the susceptibility of *C. albicans* to amphotericin B by increasing mitochondrial ROS, whilst suppressing *SOD2* expression

Superoxide radicals are removed through the actions of superoxide dismutase (Sod) enzymes. The *C. albicans* genome contains 6 SOD genes *(SOD1–6),* which vary in their subcellular localisation. In response to ROS stress, many organisms upregulate these enzymes to detoxify the superoxide radicals. However, analysis of our transcriptomic data confirmed that, in the presence of *P. aeruginosa, SOD1–5* were downregulated in *C. albicans*, while *SOD6* was upregulated (Table 1). Sod2 is a mitochondrial Sod enzyme, essential for the detoxification of mitochondrial ROS. Previous transcriptional studies confirm that, in response to oxidative stress, *SOD2* is significantly upregulated to detoxify ROS, and is usually upregulated in response to amphotericin B treatment (Liu *et al.* 2005, Wang *et al.* 2006). Given that both phenazines and amphotericin B target mitochondrial ROS production, and that menadione mimics the effects of *P. aeruginosa* on *C. albicans* susceptibility to amphotericin B, *P. aeruginosa*-induced repression of *SOD2* could enhance ROS toxicity. In agreement with this, deletion of *SOD2* resulted in significantly increased *C. albicans* susceptibility to amphotericin B in the absence of *P. aeruginosa,* while deletion of *SOD1* and *SOD3–6* had minimal effect on amphotericin B sensitivity (Fig 7A). Furthermore, addition of exogenous PMS to *sod2Δ* mono-species biofilms further enhanced the sensitivity of *C. albicans* to amphotericin B (Fig 7B). To determine whether *P. aeruginosa*-dependent suppression of *SOD2* was the driving force behind the increased sensitivity of *C. albicans* to amphotericin B, we expressed *SOD2* from a constitutive promoter and reassessed *C. albicans* viability upon amphotericin B treatment. Constitutive expression of *SOD2* resulted in increased resistance of *C. albicans* to amphotericin B in the presence of *P. aeruginosa* (Fig 7C), confirming that *P. aeruginosa-mediated* repression of *SOD2* expression contributes to the enhanced sensitivity of *C. albicans* to amphotericin B in dual-species biofilms.

**Table 1.** Expression of superoxide dismutase genes.

| ORF | Gene | Gene product | Fold change<br>(log<sub>2</sub>) | P adj |
| --- | --- | --- | --- | --- |
| **19.2060** | *SOD5* | Cu-containing Sod | -2.97 | 5.63 x10<sup>-9</sup> |
| **19.3340** | *SOD2* | Mitochondrial Mn-containing Sod | -1.85 | 5.29 x10<sup>-17</sup> |
| **19.2770.1** | *SOD1* | Cytosolic Cu- and Zn-containing Sod | -1.36 | 1.11 x10<sup>-5</sup> |
| **19.2062** | *SOD4* | Cu-containing Sod | -0.59 | 3.04 10<sup>-3</sup> |
| **19.7111.1** | *SOD3* | Cytosolic Mn-containing Sod | -0.54 | 4.10 x10<sup>-1</sup> |
| **19.6229** | *CAT1* | Catalase | 0.43 | 1.54 x10<sup>-1</sup> |
| **19.2108** | *SOD6* | Cell-surface Cu-containing Sod | 2.01 | 2.20 x10<sup>-15</sup> |

**Figure 7.**
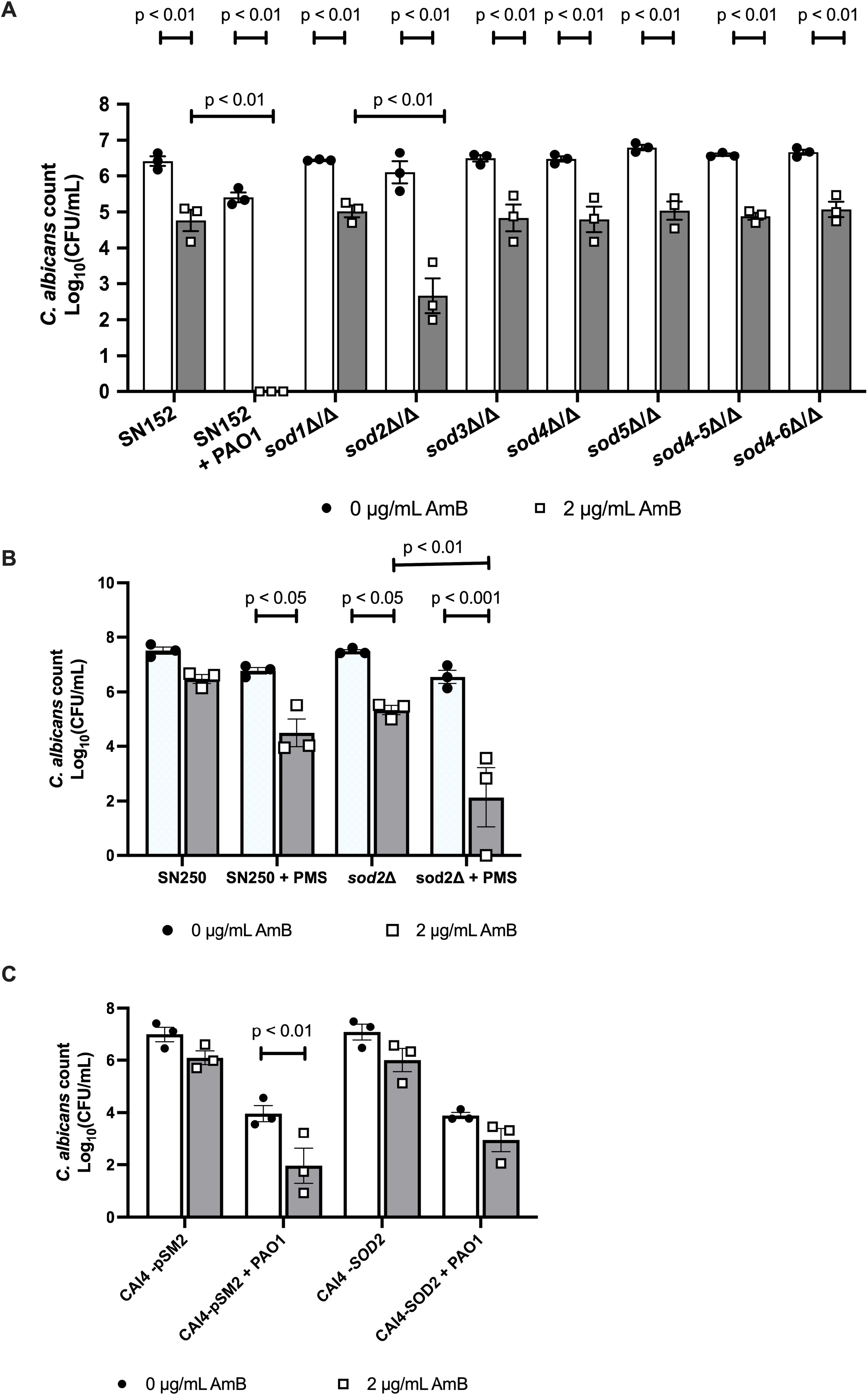
*P. aeruginosa* induces ROS stress through downregulation of the fungal mitochondrial superoxide dismutase, *SOD2*. **A)** *C. albicans* Sod mutants were grown in single-species biofilms and then treated with 2 μg/ml amphotericin B for 2 hrs. **B)** The parental control strain and the *sod2Δ* mutant were grown in single-species biofilms for 24 hrs and then treated with 2 μg/ml amphotericin B for 2 hrs. **C)** The parental control strain and the *SOD2* overexpression (CAI4-*SOD2*) strain were grown in single- or dual-species biofilms for 24 hrs and then treated with 2 μg/ml amphotericin B for 2 hrs. Data are the log_10_(mean) +/− the SEM from 3 biological replicates. Data were analysed using 2-way ANOVA and Holm-Sidak’s multiple comparisons test.

## Discussion

Antimicrobial resistance is a global threat, with multi-drug resistant strains frequently isolated in the clinic. It is now estimated that, by 2050, 10 million people per year will die as a result of antimicrobial-resistant infections (de Kraker *et al.* 2016). Although not as frequently mentioned as antibiotic resistance, antifungal resistance places a huge burden on patients and healthcare providers. This is exemplified by the isolation of multi-drug resistant strains of *C. auris*, with several strains being resistant to the three major classes of antifungals (Ostrowsky *et al.* 2020). As a result, the World Health Organisation and the Centers for Disease Control and Prevention have declared antimicrobial-resistant *Candida* infections a global health threat. Therefore, there is an urgent need for the development of new classes of antifungal drugs, or for increased efficacy of existing treatment options. Here, we have shown that *P. aeruginosa* enhances the susceptibility of *C. albicans* to amphotericin B through phenazine-dependent induction of mitochondrial ROS stress in conjunction with the suppression of ROS-detoxifying enzymes.

Owing to the nephrotoxicity of amphotericin B, it is usually used as a last-resort treatment. However, liposomal formulations have greatly reduced the toxicity of the drug. Amphotericin B targets ergosterol in the fungal membrane, impairing membrane integrity, and has been shown to induce mitochondrial ROS stress (Guirao-Abad *et al.* 2017). In agreement with this, the *C. albicans sod2Δ* mutant was more susceptible to amphotericin B than the deletion of genes encoding the other superoxide dismutases. *P. aeruginosa* has been shown to induce ROS stress in *C. albicans* through the secretion of phenazine compounds (Morales *et al.* 2010). Only PMS, a synthetic analogue of 5-methyl phenazine-1-carboxylic acid betaine (5MPCA), was able to increase the sensitivity of *C. albicans* to amphotericin B. *P. aeruginosa* secretes 5MPCA during dual-species growth with *C. albicans* (Gibson *et al.* 2009), but not during monoculture (Byng *et al.* 1979), which may explain why culture supernatants from *P. aeruginosa* monocultures did not affect the sensitivity of *C. albicans* to amphotericin B. However, supplementation of *C. albicans* mono-species biofilms with PMS did not fully recapitulate the presence of the bacterium, suggesting that additional factors are required. Supplementation of *sod2Δ* biofilms with PMS has a greater impact on the sensitivity of *C. albicans* to the polyene. *SOD2* is usually upregulated in the presence of amphotericin B and other inducers of mitochondrial ROS stress (Liu *et al.* 2005, Wang *et al.* 2006). However, transcriptional analysis confirmed that, in the presence of *P. aeruginosa*, *SOD2* and several other detoxifying enzymes were downregulated. Therefore, it would appear that *P. aeruginosa* may simultaneously induce ROS stress whilst downregulating *SOD2* expression, which likely results in the stress surpassing the capacity of the detoxification system, thus leading to cell death.

*P. aeruginosa* is highly genetically variable. Although treatment of dual-species biofilms containing PAO1 with amphotericin B resulted in minimal fungal survival after 2 hrs, biofilms containing PA14 and *P. aeruginosa* clinical isolates required a longer incubation with the drug to completely eradicate the fungus (although a significant reduction in fungal viability was observed after 2 hrs). Although all *P. aeruginosa* species produce 5MPCA, the production of the phenazine intermediate is variable between isolates (Gibson *et al.* 2009). Therefore, it is likely that, in this assay, the species produce 5PMCA at lower levels compared with the PAO1 strain, which may explain why extended incubation times are required to fully eradicate the fungus.

Other classes of antifungals also induce ROS production in *C. albicans* as part of the mode of action (Delattin *et al.* 2014), suggesting that *P. aeruginosa* should also increase the susceptibility of the fungus to other antifungals. Fluconazole targets ergosterol biosynthesis, reducing ergosterol levels in the fungal membrane, and resulting in the accumulation of toxic sterols (Yoshida 1988, Kelly *et al.* 1997). As a result, fluconazole is fungistatic against *C. albicans.* However, *in vivo P. aeruginosa* has been shown to make fluconazole fungicidal toward *C. albicans* (Hattab et al. 2022). However, in our biofilm assay, fluconazole had minimal effect on the viability of *C. albicans*, either in the absence or presence of the bacterium. However, glucans in the ECM sequester fluconazole, preventing it from penetrating the biofilm (Nett *et al.* 2010). Therefore, it is likely that, as fluconazole is added to mature biofilms, the glucan component of the ECM sequesters the drug, preventing it from reaching the fungal cells.

Although *C. albicans* is still the main species causing candidiasis, the incidence of infections involving non-*albicans Candida* species is increasing (Giacobbe *et al.* 2020). Therefore, it is important for antifungal therapies to be effective across all *Candida* species. *P. aeruginosa* enhanced the susceptibility of *C. dubliniensis* to amphotericin B, similarly to *C. albicans*, which is not surprising, as *C. dubliniensis* is the most closely related species to *C. albicans*. Amphotericin B has a negligible effect on the viability of *C. krusei* in the presence or absence of *P. aeruginosa*, suggesting that this isolate is resistant to the polyene. Estimates suggest that up to 15% of *C. krusei* isolates are resistant to amphotericin B owing to reduced ergosterol content in the cell membrane (Krcmery and Barnes 2002). Despite the other *Candida* species being susceptible to the polyene, their susceptibility was not significantly enhanced in the presence of *P. aeruginosa.* The interactions between *P. aeruginosa* and non-*albicans Candida* species have not been as extensively studied as *C. albicans*. Therefore, it is possible that either 5MPCA does not induce ROS stress, that the bacterium is not able to downregulate the ROS detoxifying enzymes in these species, or that these species can tolerate greater levels of ROS than *C. albicans.*

In the clinical setting approximately 80% of infections are due to the formation of biofilms, which are highly antimicrobial resistant due to the upregulation of efflux pumps and the extracellular matrix providing protection from the environment (Van Acker *et al.* 2014, Lohse *et al.* 2018, Rodrigues *et al.* 2020). Therefore, *in vivo*, much higher concentrations of drug are required to eradicate the infection, but cannot always be achieved. Our data confirm that, in a mixed-species biofilm, *C. albicans* is more susceptible to amphotericin B than mono-species biofilms. Given that *in vivo* biofilms are normally composed of multiple species, especially in traumatic wounds and in the CF lung where *C. albicans* and *P. aeruginosa* are frequently co-isolated, this suggests that *C. albicans* will be more susceptible to amphotericin B than the other classes of antifungal drugs. Furthermore, we and others have shown that *C. albicans* promotes drug tolerance in multiple bacterial species (Harriott and Noverr 2009, Kong *et al.* 2016, Alam *et al.* 2020). Therefore, *Pseudomonas* biofilm infections that contain *C. albicans* may be best treated with an initial course of antifungal, prior to the initiation of antibacterial treatment.

## Acknowledgements

Work in the Hall Lab is supported by an MRC Career Development Award (MR/L00903X/1) and the BBSRC (BB/R00966X/1, BB/R00966X/2). Work in the Blair Lab is supported via a David Phillips Fellowship to J.M.A.B. (BB/M02623X/1). F.A. is supported by the Wellcome Trust Antimicrobials and Antimicrobial Resistance (AAMR) doctoral training programme (108876/Z/15/Z).

## Author contributions

RAH and JB conceived and designed the experiments, FA, SB and JC performed the experiments and analysed the data. RAH wrote the manuscript.

---

**Figure S1. Transcriptional profiling of *C. albicans* in single- and dual-species biofilms. A)** PCA plot. **B)** Volcano plot of significantly differentially regulated genes (p <0.05 and log_2_ fold change > 1).

